# Pre-conscious reactions to faces as the biological roots of emotion perception

**DOI:** 10.1101/2025.08.19.671017

**Authors:** Cecilia Dapor, Ronen Hershman, Daniela Ruzzante, Federica Meconi, Irene Sperandio

## Abstract

Bodies respond to others’ emotions through subtle physiological changes. Whether these responses require conscious emotion recognition is debated. Here, we examined the relationship between visual awareness and physiological arousal in response to emotional faces. We recorded facial electromyography (EMG), electrodermal activity (EDA), and pupil dilation during the presentation of fearful, happy, and neutral faces, as well as their phase-scrambled versions, to the left eye. Meanwhile, rapidly changing Mondrian patterns were displayed to the right eye to suppress the left-eye stimuli. Participants pressed a button when they noticed a change from the flickering mask. This setup allowed us to compare behavioural and physiological responses during conscious versus subconscious processing of emotional content. Results showed faster breakthrough times for happy faces and moderate physiological responses to emotional information, even without conscious perception. Specifically, pupils and facial EMG were more responsive to fearful than to happy or neutral expressions. EDA was higher for faces compared to control stimuli, regardless of the emotional expression. Once visual awareness was established, emotional expressions elicited distinct responses that aligned with the perceived emotion. An exploratory analysis revealed a negative correlation between pupil dilation and autistic, alexithymic, and schizotypal traits, whereas empathy traits correlated positively. Our findings highlight the role of subcortical visual processing in detecting emotionally relevant stimuli and show that social abilities are linked to the physiological processing of emotions at a subconscious level. They underscore how subcortical mechanisms interact with the autonomic nervous system, enhancing our understanding of the biological foundations of emotion perception.

## Introduction

To navigate our social world, we must rapidly interpret others’ emotions, such as recognizing an angry colleague or a fearful passerby, and prepare appropriate responses. A brief glance at a face or body can reveal another person’s psychological state [1], triggering an autonomic reaction in our own body. These embodied responses facilitate emotion understanding and support adaptive social behaviour. Yet, the unconscious physiological mechanisms that enable this rapid social attunement remain poorly understood. In particular, it is still debated whether autonomic responses arise before conscious emotion recognition, and to what extent these early bodily reactions contribute to social cognition.

Research in affective neuroscience suggests that emotional facial expressions benefit from specialized neural mechanisms. A key component is the subcortical visual pathway, which rapidly transmits visual information to the amygdala and other subcortical structures, supporting fast, reflexive responses. Indeed, neuroimaging studies have shown amygdala activation to emotional faces even when presented subliminally [2–5], suggesting that a primitive form of emotion discrimination can occur unconsciously. The amygdala also plays a central role in autonomic regulation, allowing physiological reactions to emotional cues before we become aware of them.

Further evidence in this direction comes from the work of Celeghin and colleagues [6] on patients with lesions in the primary visual cortex. They were able to recognize the emotional expressions presented to the blind hemifield above the chance level and exhibited facial mimicry, despite lacking conscious awareness of the stimuli. These findings indicate that emotional stimuli can trigger bodily responses even in the absence of visual awareness. Beyond facial mimicry, other autonomic indices, including electrodermal activity (EDA), heart rate variability, skin temperature, and pupil dilation, can reveal the physiological expression of emotion perception [7]. Pupil dilation, for instance, reliably reflects sympathetic arousal [8] and previous research indicates that fearful facial expressions elicit greater pupil dilation than happy or neutral ones, even when the stimuli are presented subliminally [9]. However, most studies have examined these physiological markers in isolation, limiting our understanding of how different components of the autonomic nervous system jointly contribute to unconscious emotional processing.

In healthy individuals, the unconscious processing of facial expressions has been widely investigated through masking paradigms and interocular suppression paradigms [10,11]. Standard backward masking and object-substitution masking can render stimuli invisible for up to ∼100 ms, but these brief intervals limit their utility for autonomic research. Continuous Flash Suppression (CFS) overcomes this limitation by presenting a dynamic series of high-contrast “Mondrian” images to one eye while the target stimulus is shown to the other, inducing interocular rivalry that can suppress the target from awareness for several seconds [12,13]. This prolonged suppression has made CFS a robust tool for probing preconscious processing and its neural correlates. However, its effectiveness at fully suppressing visual processing is debated [14–16], and its reliance on subjective reports limits its objectivity.

A widely used variant, breaking Continuous Flash Suppression (bCFS) [17], measures the time it takes for a suppressed stimulus to “break through” to awareness. Stimuli that overcome suppression faster are assumed to benefit from prioritized unconscious processing. Numerous studies have reported that fearful facial expressions break through suppression faster than other expressions, suggesting an advantage for threat-related cues [18,19].

However, subsequent research has challenged this interpretation, demonstrating that differences in low-level visual features, such as local contrast, or spatial frequency, can significantly influence breakthrough times [14,20]. Moreover, bCFS reaction times cannot clearly distinguish between stimulus detection and identification [14], introducing a strong confound on the boundary between unconscious processing and conscious access.

Despite its limitations, the widespread use of the b-CFS technique and the substantial body of research it has generated make it a strong candidate for exploring emerging topics such as the preconscious physiological processing of emotions. In the present study, we combined bCFS with simultaneous measures of EDA, pupillary responses, and facial electromyography to investigate whether emotional facial expressions evoke distinct physiological responses before reaching awareness. Participants responded as soon as the stimulus broke through suppression, then identified the perceived emotion and provided a subjective clarity rating of their visual experience using a four-point version of the PAS [21]. To ensure that the pre-response physiological signals reflected genuinely pre-conscious processing, we anchored their analysis to stimulus onset rather than to the behavioral response. For each physiological measure, we defined time windows based on its minimal modulation latency, capturing responses elicited precisely at stimulus appearance, when visual awareness suppression should be maximal. Furthermore, to better dissociate detection from identification effects on the response time, we run a complementary follow-up experiment using the same technique without requiring explicit emotion recognition. This multimodal approach allows for a more comprehensive assessment of autonomic activation during unconscious processing and enables direct comparison of different physiological indices. We also explored whether preconscious responses are increased in individuals with high social-affective traits, including empathy, extending prior work that has linked atypical autonomic reactivity to clinical and subclinical changes in social functioning [22–24]. Indeed, prior research suggested that individuals with higher levels of autistic, schizotypal, or alexithymic traits may exhibit reduced physiological activation in response to emotional faces. Conversely, we expected that higher empathy scores would be associated with increased pre-conscious physiological reactivity.

Our findings demonstrate that emotional faces, particularly fearful ones, elicit measurable physiological responses prior to conscious recognition. Moreover, pupillary responses to unseen emotional faces correlated negatively with autistic, alexithymic, and schizotypal traits, and positively with empathy. Together, these results indicate that the human body resonates with emotional cues even in the absence of awareness, and that variability in this preconscious resonance is higher for individuals with high social-affective functioning.

## Results

### Behavioural findings: Breakthrough times and accuracy

Timing differences in breakthrough suppression across different stimulus categories depend on their relevance and priority for our visual system. Stimuli that are detected faster than others afford preferential processing due to their relevance for our immediate behavioural outcomes [25]. Social stimuli fall into this special category, as confirmed by our reaction time (RT) data (see Figure 1a). Results from the 2×3 repeated-measures ANOVA with Image Type (faces vs. scrambled) and Emotion (fearful vs. happy vs. neutral) as main factors revealed that participants’ mean RTs for faces were significantly faster than those for the scrambled images (i.e., a main effect of Image Type: *F*(1, 66) = 19.53, *p* < .001, *ηp^2^* = .228). No effect on RTs was found for Emotion (*F*(2,132) = 2.885, *p* = .06). However, the interaction between Image type and Emotion was significant, *F*(2,132) = 11.278, *p* < .001, *ηp^2^* = .146, indicating that the effect of the emotional content was specific to the faces.

**Figure 1.**
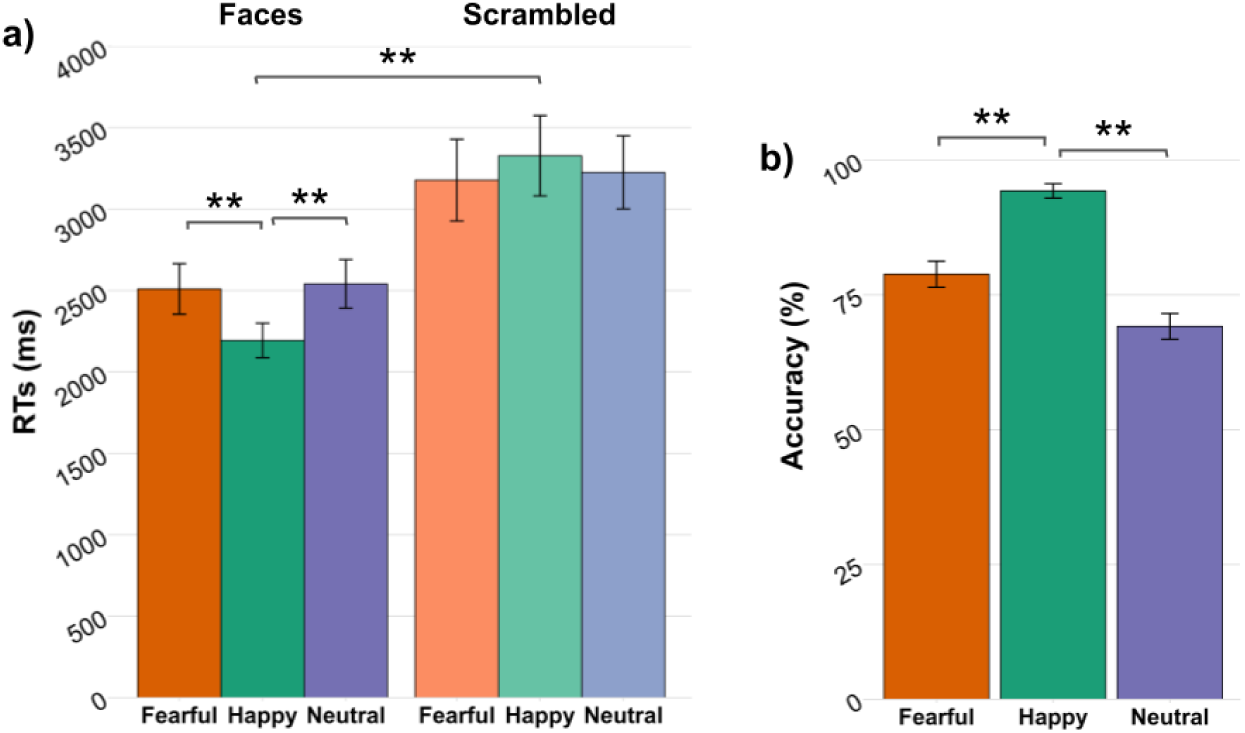
Mean breakthrough times for all the conditions **(a)** and accuracy percentages for faces **(b)**. Error bars correspond to the standard error of the mean. Asterisks denote significant differences at p < .05 (*) and p < .01 (**).

Post-hoc analysis confirmed that happy faces were detected faster than fearful (*t*(66) = 3.767, *p_bonf_* < .001, *d* = 0.464) and neutral ones (*t*(66) = 4.514, *p_bonf_* < .001, *d* = 0.556), whereas no effect of emotion was found for their scrambled counterparts (Figure 1a).

The one-way repeated-measures ANOVA on the emotion recognition of facial expressions revealed significant differences in accuracy driven by the specific emotion (*F*(2, 130) = 46.875, *p* < .001, *ηp^2^*= .419). Post-hoc t-tests confirmed that happiness was the most accurately recognized emotion, followed by fearful and neutral expressions (Figure 1b). There was a significant difference in accuracy between recognizing happy expressions and both fearful (*t*(65) = 6.43, *p_bonf_* < .001, *d* = .79) and neutral (*t*(65) = 11.163, *p_bonf_* < .001, *d* = 1.37) expressions. Furthermore, fearful expressions were recognized more accurately than neutral ones (*t*(66)= 3.221, *p_bonf_* = .006, *d* = .4).

### Physiological findings: Electrodermal activity, facial EMG, and pupillometry

To determine whether emotional faces elicit physiological reactions before reaching conscious awareness, we measured EDA, facial EMG, and pupil dilation in time windows both preceding and following participants’ responses. Because visual masking paradigms raise ongoing debate about the precise moment a stimulus enters consciousness [14,26,27], we addressed this challenge by calibrating our analysis windows to the known latencies of each physiological measure. In particular, to isolate the unconscious response, we selected the earliest time window where a reaction linked to the stimulus onset was physiologically possible. A corresponding post-awareness window was defined relative to the participants’ response. In this way, we obtained two values for all the compared conditions: a pre- and a post-awareness response. Accordingly, a third within-subject factor (Awareness: pre vs. post) was included in the repeated-measures ANOVA used for the EDA and EMG analysis. Pupil dilation was analysed following a different approach, specifically through a series of Bayesian t-test comparisons on the time-course data (following Hershman and colleagues [28]). Trial accuracy was not included in the analyses because error rates were minimal, likely reflecting the low difficulty of the recognition task, and their distribution across conditions was uneven. Similarly, PAS scores showed a highly skewed distribution, which made them less suitable for trial-by-trial analyses. Instead, they were used in correlational analyses with the physiological data to control for potential effects of perceptual awareness on pre-conscious responses (see Supplementary Materials for all analyses and results). Further details on participant inclusion, power analyses, and exclusion criteria are provided in the method section (available online).

### EDA

EDA was analyzed using the mean Integrated Skin Conductance Response (ISCR) extracted from a two-seconds time window, either starting one second after stimulus onset (pre-awareness response) or following the participant’s response (post-awareness response). Event-related EDA is a relatively slow physiological signal, typically emerging about one second after stimulation [29]. By restricting the time window within three seconds after stimulus onset and considering that the average RT exceeded two seconds, we minimized any potential overlap with the EDA elicited by conscious perception, which was calculated in the 1–3 second window after the participant’s response. The three-way repeated-measures ANOVA revealed a main effect of Image type (*F*(1,50) = 6.483, *p* = .01, *ηp^2^* = .115), indicating that the social stimuli (i.e. faces) elicited greater EDA than their scrambled counterparts (Figure 2). Notably, there was no effect of Awareness (*F*(1,50) = .871, *p* = .35, *ηp^2^* = .017), indicating that differences in EDA occurred independently of participants’ visual awareness of the stimuli. The remaining main effect (Emotion) and interactions did not reach significance (*max F* = 2.788, *min p* = .07). This result suggests that social stimuli can induce an electrodermal response whether or not the stimulus is consciously perceived.

**Figure 2.**
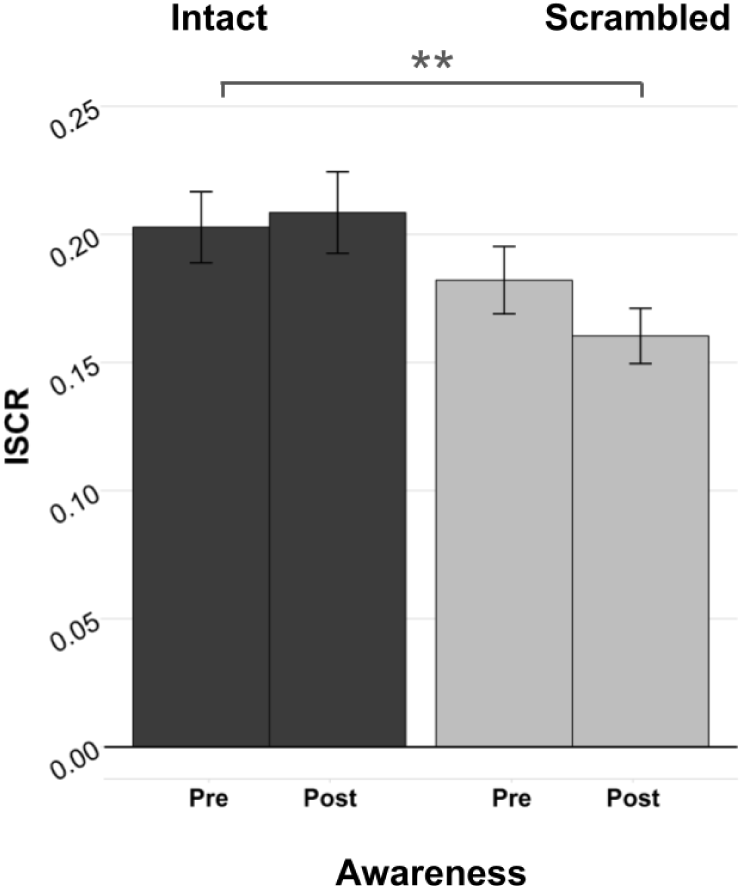
Mean integrated skin conductance response (a measure of EDA) to intact faces (left) and their scrambled versions (right), pre and post-visual awareness. Error bars correspond to the standard error of the mean. Asterisks denote significant differences at p < .05 (*) and p < .01 (**).

### EMG

The EMG activity of the zygomaticus and the corrugator muscles was analyzed separately. Given that facial mimicry typically emerges within 500 ms after stimulus presentation [30], we focused on this time window to extract the EMG activity. We calculated the mean EMG response for the first 500 ms bin after stimulus onset, corresponding to the pre-conscious period, and the first 500 ms after the participant’s response, corresponding to the conscious period. The analysis of the corrugator supercilii response did not yield any significant results (no main effects nor interactions, *max F* = 1.776, *min p* = .19). A possible explanation for this lack of findings could be that the corrugator supercilii shows a frowning-related activity that is often associated with a general state of focus, rather than facial mimicry [31].

The three-way repeated-measures ANOVA of the zygomaticus major response revealed a significant main effect of Awareness (*F*(1,54)= 6.574, *p* = .01, *ηp^2^* = .109), indicating greater muscle activity after the stimulus reached conscious awareness. The interactions of Awareness with the other main factors were not significant (*max F* = 1.46, *min p* = .23). Neither the main effect of Emotion (*F*(2,108)= 1.936, *p* = .14) nor the main effect of Image Type (*F*(1,54)= 2.725, *p* = .10) reached significance. However, there was a significant interaction between Image Type and Emotion (*F*(2,108) = 6.321, *p* < .01, *ηp^2^* = .105), which demonstrates emotion-specific muscle activation even during the unconscious processing of the faces. In particular, fearful faces elicited greater activation than neutral expressions already in the pre-conscious phase (*t*(54) = 2.734, *p_bonf_* = .024, *d* = .35). Zygomaticus activity was also greater for happy faces compared to neutral ones; however, this difference did not remain statistically significant after applying the Bonferroni correction (*t*(54) = 2.119, *p_uncorr_* = .038, *p_bonf_* = .114, *d* = .27). The atypical activation of the zygomaticus muscle in response to fearful expressions might reflect motor resonance rather than mimicry, as the fearful faces in the study displayed a widely opened mouth that naturally engages the muscles surrounding the zygomaticus. A different pattern of activation was observed after the conscious appraisal of the stimuli, as confirmed by the significant three-way interaction between Awareness, Image Type, and Emotion (*F*(2,108) = 3.783, *p* = .03, *ηp^2^* = .06). Happy facial expressions elicited significantly greater activation of the zygomaticus major compared to neutral expressions (*t*(54) = 3.763, *p_bonf_*< .001, *d* = .37). Activation was also greater in response to happy faces than to fearful ones, although this difference did not reach significance after Bonferroni correction (*t*(54) = 2.102, *p_uncorr_* = .04, *p_bonf_* = .12, *d* = .28). The results indicate the presence of motor resonance when the facial expression is masked. Interestingly, fearful expressions are mirrored when unconsciously perceived, while after reaching visual awareness, the zygomaticus activation is significantly greater only for happy expressions, as is usually found in the literature. Figure 3 represents the zygomaticus activation for faces both before and after visual awareness. Importantly, the scrambled images had no effect on the muscles (all p-values > .05), confirming the social drive of facial motor resonance.

**Figure 3.**
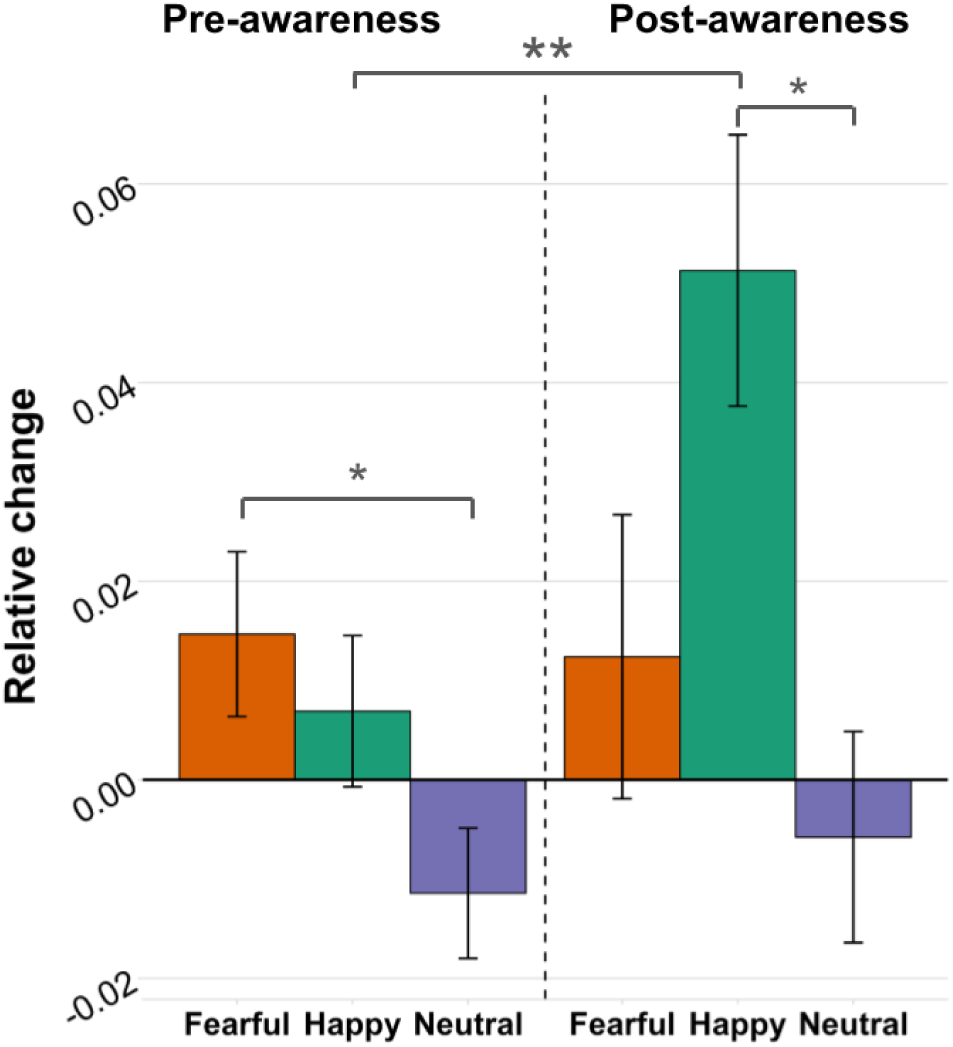
Mean relative change (compared to baseline) of zygomaticus major activity for the three emotional faces, before (left) and after (right) visual awareness. Error bars correspond to the standard error of the mean. Asterisks denote significant differences at p < .05 (*) and p < .01 (**).

### Pupil dilation

To analyze differences in pupil dilation across conditions, we conducted a series of Bayesian paired-samples t-tests over the averaged time-course of two different epochs (i.e., stimulus-locked and response-locked), corresponding to pre- and post-visual awareness levels. We first compared the mean dilation for faces versus scrambled images, and then we separately tested the three emotions in both the intact faces and their scrambled versions.

Figure 4 shows that there was a significant difference between intact and scrambled images only after the conscious perception of the stimuli. More specifically, the difference between the two conditions became significant ∼200 ms after the participant’s response and remained significant for the entire duration of the post-stimulus fixation cross (4000 ms, Figure 4b). No significant difference was observed in the time window locked to the onset of the stimuli (Figure 4a).

**Figure 4.**
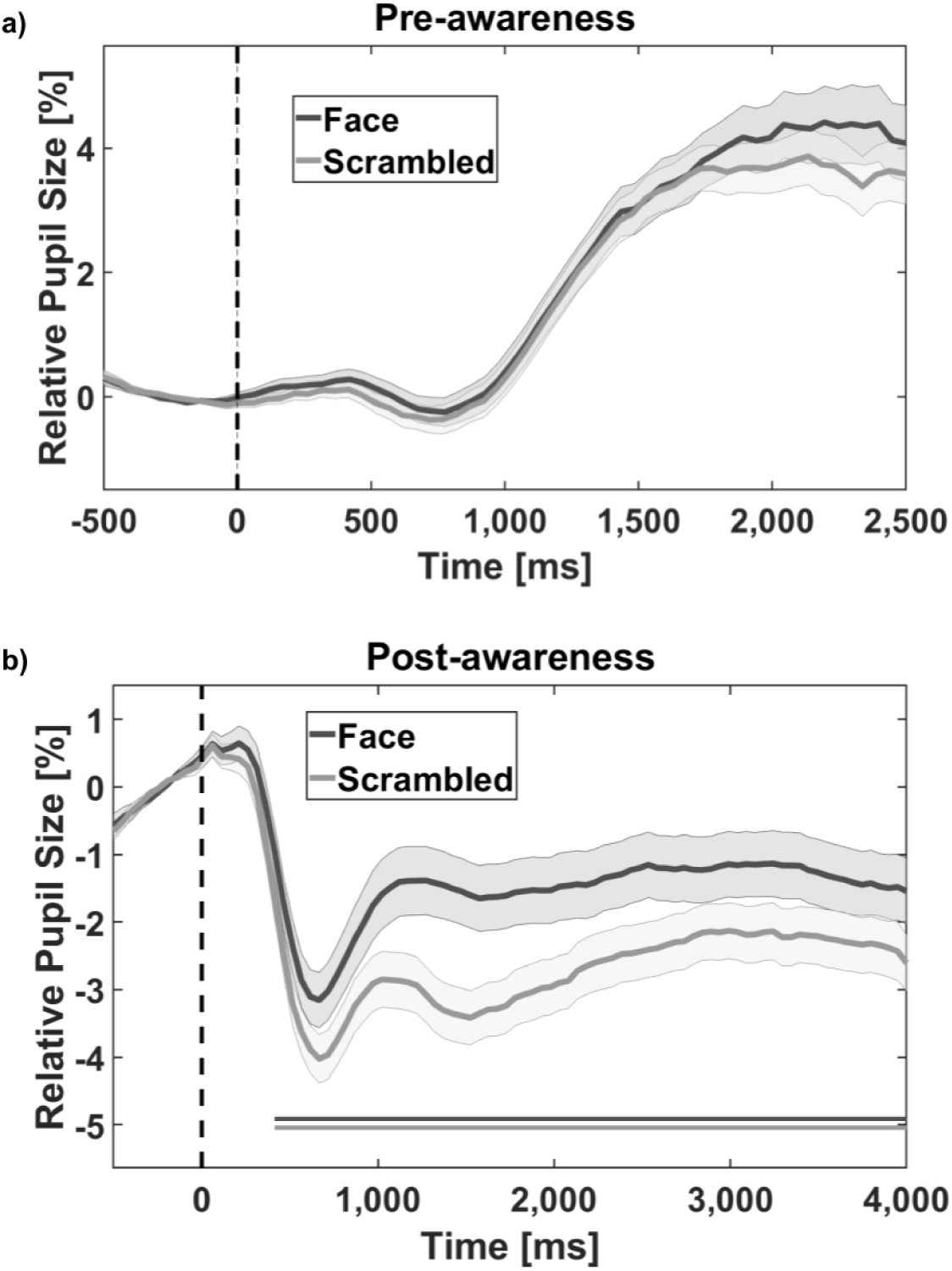
Mean relative pupil size (with respect to a pre-stimulus baseline of 500 ms) for the two image types (intact vs. scrambled), before **(a)** and after **(b)** visual awareness. The dashed vertical line at time-point zero represents stimulus onset in a) and response time in b). The line curves represent changes in pupil dilation as a function of time. The shaded areas represent one standard error from the mean. Solid horizontal lines under the graphs indicate significant differences between conditions (i.e., *BF*_10_ ≥ 3).

Figure 5 illustrates pupil changes in response to the processing of emotional information. We registered meaningful differences (i.e., BF_10_ ≥ 3) in pupil change around 2,100 ms after stimulus onset: pupils dilated more in response to fearful than to happy faces and remained sustained until the participant’s response, before the stimuli entered conscious awareness (Figure 5a). Meaningful differences related to facial expressions were also observed later when stimuli were consciously perceived (Figure 5b). Fearful faces elicited greater dilation than happy faces, starting at approximately 1,200 ms after the breakthrough, and the difference remained significant for 1,000 ms.

**Figure 5.**
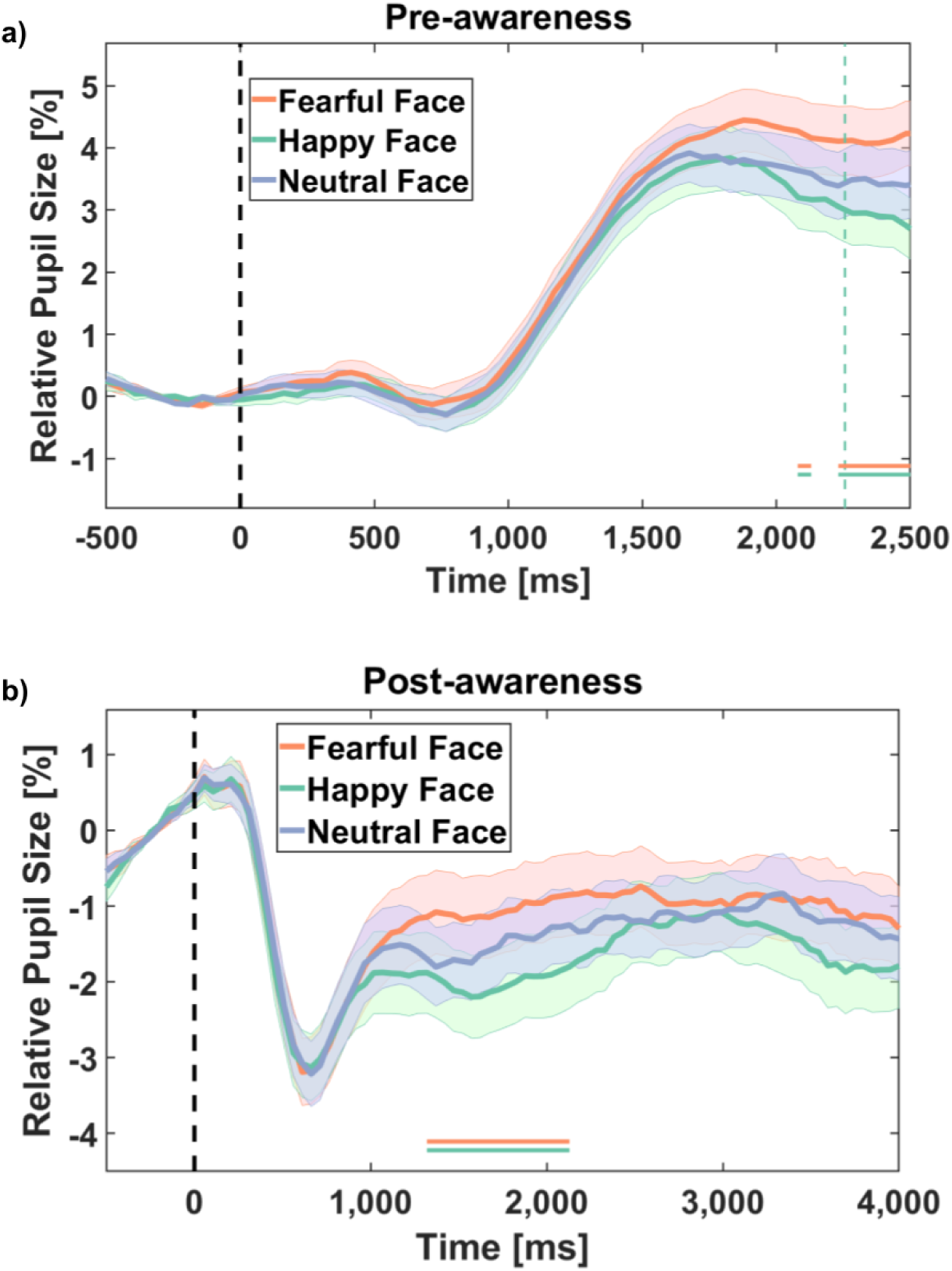
Mean relative pupil size (with respect to a pre-response baseline) for the three emotional faces before **(a)** and after **(b)** visual awareness. The dashed vertical line at time-point zero represents stimulus onset in a) and response time in b). The shaded areas represent one standard error from the mean. Solid horizontal lines under the graphs indicate significant differences between conditions (i.e., BF_10_ ≥ 3).

Importantly, none of the comparisons between the different scrambled conditions yielded any significant results, either before or after visual awareness. Taken together, the findings indicate that pupil dilation is modulated by emotional content, even in the absence of visual awareness. In particular, fearful expressions elicited the greatest dilation both before and after the conscious perception of the facial expressions.

### Interindividual differences in physiological arousal

To examine whether individual dispositional traits influence perceptual sensitivity to social stimuli, we calculated correlations between self-report measures (autistic traits, empathy, negative emotional states, alexithymia, and schizotypy) and the physiological responses measured in the pre-conscious time window. We expected physiological responses to be positively correlated with social abilities and negatively correlated with social-affective difficulties. To reduce the number of variables, we computed a single sensitivity index for social stimuli by subtracting the mean response to scrambled images from the mean response to intact faces, obtaining two values for each physiological index, one before and one after visual awareness.

The results revealed significant correlations between individual differences in personality traits as well as emotional states and the pupillary response during the pre-conscious phase. Specifically, we found a negative relationship between changes in pupil diameter and AQ (*r*(40) = −.353, *p_uncorr_* = .011), SPQ (*r*(39) = −.308, *p_uncorr_* = .025), TAS (*r*(40) = −.348, *p_uncorr_* = .012), and the Depression subscale of the DASS (*r*(39) = −.338, *p_uncorr_* = .015) scores. In contrast, EQ correlated positively with pupil size (*r*(40) = .289, *p_uncorr_* = .03). Therefore, our findings showed that increased levels of social impairments (AQ, SPQ, TAS) or emotional distress (DASS) were associated with decreased pupil reactivity to faces, while empathy (EQ) exhibited the opposite trend: the higher the empathic abilities, the greater the dilation of the pupil. None of the remaining correlations reached significance. It should be noted that these relationships were observed only in the pupillometry data and not in the other physiological indices. Graphs of all correlations in the pre-conscious phase are represented in Figure 6. Results from the correlational analysis of the post-conscious phase are reported in the Supplementary Materials.

**Figure 6.**
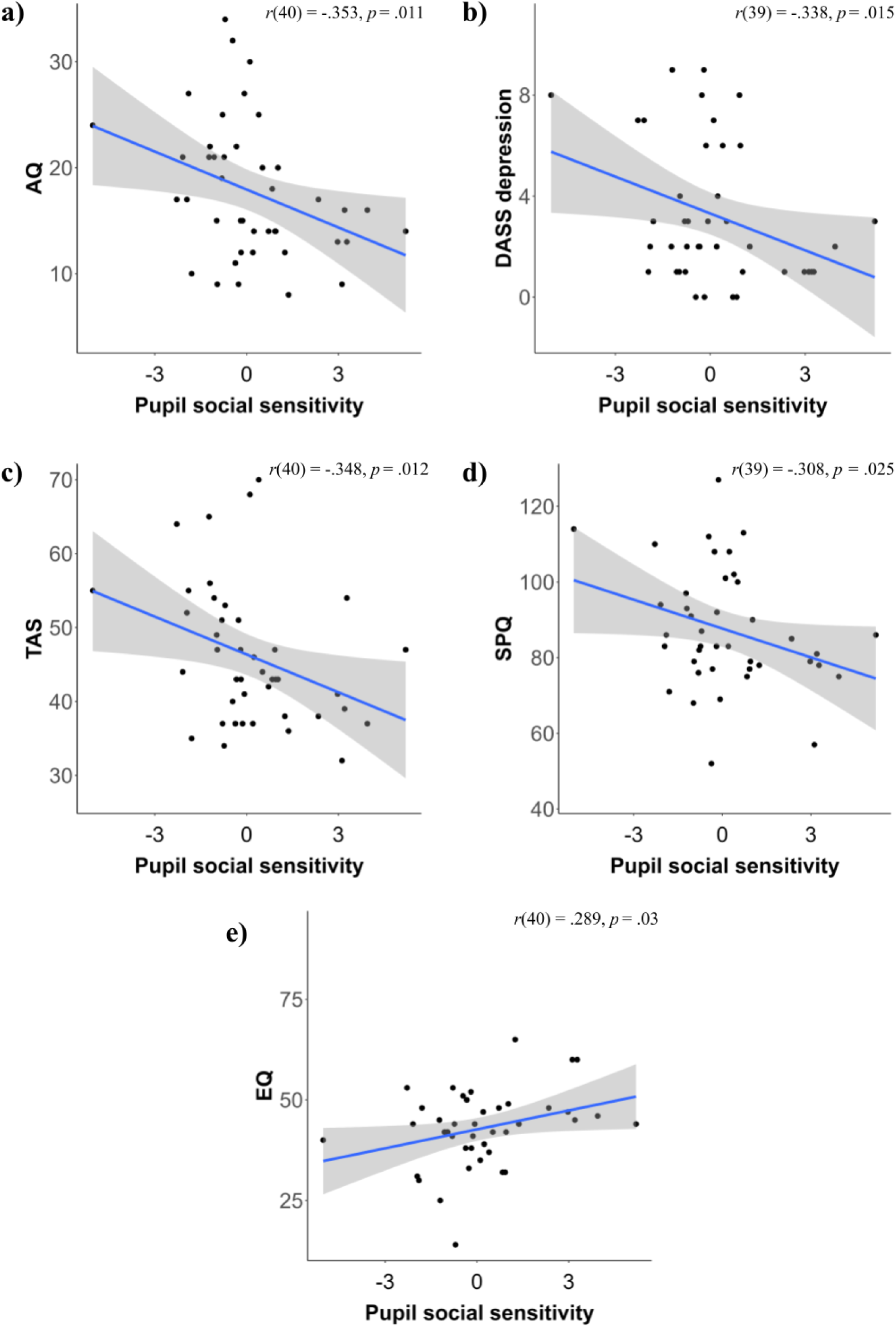
Correlations between pre-conscious pupil dilation and personality questionnaires. Positive values on the pupil social sensitivity axis indicate greater dilation toward faces compared to scrambled images. The graphs show the positive correlations between pre-conscious pupil dilation and AQ (a), the DASS Depression subscale (b), TAS (c), and SPQ (d). The graph at the bottom (e) shows the negative correlation between pupil dilation and EQ.

## Discussion

In this study, we aimed to uncover the psychophysiological mechanisms underlying emotion detection and recognition, focusing on the automatic and pre-conscious processing of facial expressions. Using the bCFS paradigm and autonomic activation measures, we examined both pre-conscious and conscious responses to emotional faces and observed an emotion-specific pattern of physiological activation even before participants became consciously aware of the stimuli. These findings build upon previous research on the subcortical processing of emotional stimuli, which can activate the autonomic nervous system independently of visual awareness, especially thanks to the amygdala’s involvement [32–34]. Furthermore, we showed that individual social skills modulate the pupillary response. Pupil dilation is a reliable measure of attentional processes and cognitive load, and it also reflects autonomic activation linked to states of arousal (e.g., during emotion recognition) [8,35]. Among the physiological measures we recorded, pupil dilation proved to be the most sensitive indicator of differences in cognitive styles. Specifically, individuals with greater dispositional social resources exhibited a pre-conscious modulation of pupil diameter, indexing an individual predisposition to detect social stimuli.

In the ensuing discussion, we will first report our behavioural and physiological findings and then discuss how individual differences in personality traits and emotional states relate to the different physiological indices.

### Is the privileged detection of emotional faces linked to task demands?

We observed faster breakthrough times for faces compared to their scrambled counterparts and a privileged access to consciousness for happy faces relative to fearful and neutral expressions. This preferential access to awareness of faces replicates previous findings and is considered a hallmark of visual masking studies [10,36]. However, reaction times in our primary experiment are likely influenced by the emotion identification process, which can delay participants’ responses [14]. The literature on emotion-specific processing has produced mixed findings, and our results do not align with several bCFS studies reporting that fearful expressions break suppression more rapidly than other emotions [18,20]. Importantly, most of these studies did not include an explicit emotion recognition component. To examine potential task-related effects, we conducted a follow-up experiment with a smaller sample in which the emotion recognition task was replaced by a simple detection task (for more details, see the Supplementary Materials). This control experiment revealed that faces break through suppression independently of their emotional content: although faces were detected faster than their scrambled counterparts, the advantage for happy faces disappeared. This suggests that the effect observed in our main experiment was likely driven by the inclusion of the recognition task. Consistent with Becker and colleagues [37], the apparent superiority of happy faces may reflect the readily interpretable communicative intent of a smile, whereas fearful and neutral expressions are more ambiguous and slower to categorize, particularly when they must be distinguished from similar emotions such as surprise or sadness.

### Physiological findings

Each physiological index captures a distinct aspect of autonomic nervous system activity. As Folz and colleagues [7] note, sympathetic activation is not an “all-or-none” response but instead involves complementary contributions from different physiological systems. As such, we have examined the specific patterns of physiological activation exhibited by each index during the unconscious processing of visual stimuli. Importantly, we ruled out the possibility that the pre-conscious physiological reactions were influenced by different degrees of perceptual awareness by examining the effects of the Perceptual Awareness Scale (PAS) on each physiological response through correlational analysis (see Supplementary Materials).

Electrodermal activity was reliably enhanced for faces compared to scrambled images both before and after awareness, indicating that the autonomic system distinguishes socially relevant from irrelevant stimuli without conscious recognition. This result aligns with a substantial body of evidence linking amygdala activation to increased EDA in response to expressive faces [38–41]. However, EDA was not modulated by emotional content, likely due to the low arousal elicited by static facial images, which do not convey an immediate threat, and the simplicity of the task, which may have limited participants’ emotional engagement.

Facial EMG revealed a selective modulation of the zygomaticus major muscle. Before awareness, fearful faces elicited the greatest activation, followed by happy faces. This unexpected response to fear likely reflects stimulus-specific features, as our fearful expressions included a wide mouth opening that may have engaged zygomaticus-adjacent muscles via motor resonance rather than pure emotional mimicry. After conscious detection, the response of the zygomaticus major to fear diminished, and its activity became primarily driven by happy faces. This finding aligns well with the literature on facial mimicry, which posits that people tend to automatically mimic the facial expressions of others as a component of social communication and empathy [2,42,43]. Corrugator supercilii activity showed no reliable modulation, consistent with its association with cognitive effort rather than specific negative mimicry. These findings suggest that automatic, pre-conscious facial responses can emerge independently of visual awareness, supporting evidence from TMS [44] and masking studies [45] of early motor resonance to emotional faces.

Pupillometry provided convergent evidence of pre-conscious activation of the autonomic nervous system in response to faces. Although initial differences between faces and scrambled images were subtle, faces evoked stronger dilation shortly after breakthrough, consistent with pupillary responses to emotional content emerging with a ∼2 s latency [8,46–48]. Critically, fearful faces elicited larger pupil dilations than happy faces even before awareness, reflecting sympathetic dominance in threat processing and replicating prior findings of subcortical pupillary responses to fear [9,49].

Together, these convergent physiological measures support the view that faces, and in particular threatening expressions, trigger rapid, automatic autonomic and motor responses, some of which emerge even before visual awareness.

### Interindividual differences in pre-conscious physiological modulation: Implications for emotional processing

An innovative aspect of our study was the investigation of how personality traits modulate pre-conscious physiological responses to emotional faces. While individual differences in conscious perceptual and physiological processing have been extensively investigated, especially in clinical populations, less is known about how these differences manifest pre-consciously. Prior evidence shows atypical autonomic reactivity in autism and schizophrenia, suggesting that physiological differences are closely linked to social functioning [22,23,50,51]. Drawing on embodied social cognition theories [52,53], we hypothesized that higher social predisposition (i.e., higher levels of empathy indexed by the EQ) would be associated with stronger pre-conscious physiological responses to faces, whereas higher social difficulties (i.e., higher levels of autistic, schizotypal, and alexithymic traits indexed by the AQ, SPQ, TAS, respectively) would exhibit the opposite pattern.

This hypothesis was supported by the findings on the pupillary response. Pre-conscious pupil dilation correlated positively with EQ scores and negatively with autistic, schizotypal, and alexithymic traits, indicating that social disposition influences autonomic reactivity even before awareness. Interestingly, higher depressive symptoms (DASS) were also associated with reduced pre-conscious pupil dilation, consistent with blunted emotional engagement or diminished attention to social cues [54]. These results align with reports of pupillary anomalies in autism and alexithymia [55,56], and with evidence of pupillometry sensitivity to perceptual differences associated with autistic traits in the non-clinical population [57]. Similarly, alexithymia has been associated with autonomic hyporeactivity [24], which is in line with our findings. It is important to note that our experiment required a relatively low cognitive load, as the emotion recognition task was rather easy. Experiments involving more complex social abilities may reveal a different pattern, with individuals experiencing social difficulties (e.g., those with autism) displaying greater pupil dilation due to the increased cognitive load required by the task, as demonstrated by Lee and colleagues [58].

To our knowledge, this is the first study to indicate that interindividual differences in personality and affective states shape physiological activation at the earliest stages of emotion processing, even in a non-clinical population. Although our sample size imposes some limitations and may have reduced statistical power of the correlational analyses, these findings highlight a promising avenue for future research, particularly in linking early physiological resonance to social predispositions and broader individual differences in socio-emotional functioning.

### Debate on CFS

The bCFS paradigm we employed provides a valuable tool for investigating pre-conscious processing; however, its interpretability is constrained by several well-recognised limitations [14,59]. One common critique concerns the persistence of low-level image features that may facilitate visual processing [16,20]. To mitigate this, we carefully measured and matched key low-level stimulus properties, including luminance and spatial frequency, while acknowledging that residual differences in other features cannot be fully excluded. Our primary interest, however, was in physiological responses driven by the stimuli’s emotional content, which in our data consistently matched the displayed emotional category. Such emotion-specific patterns are less likely to be explained by low-level image properties, which tend to be more influential in determining breakthrough times, thereby reducing the relevance of these potential confounds for our analyses.

The PAS, developed to quantify levels of perceptual awareness under the assumption that consciousness is a graded phenomenon [21], is often used alongside CFS tasks to provide a complementary, subjective measure of awareness. In our study, however, the PAS showed a highly skewed distribution, with near-ceiling values for faces, likely due to the combination of self-paced responses and a recognition task. Consequently, we could only conduct correlational analyses to control for potential effects on pre-conscious physiological signals. This limitation was partially addressed by calibrating the time window for physiological analysis as outlined above.

Future research would benefit from incorporating stronger and less biased measures of awareness, such as paradigms that separate detection from discrimination and yield bias-free sensitivity estimates (e.g. [11,27,60,61], to more rigorously disentangle conscious from unconscious processing. Despite these methodological constraints, we remain confident that our findings are robust and represent a solid foundation for further research integrating physiological indices into the study of visual awareness.

## Conclusions

Our study provides a comprehensive view of how visual awareness interacts with autonomic and motor responses by combining behavioural measures with multiple physiological indices.

Notably, in our experiment, pupillometry emerged as the most sensitive measure of pre-conscious emotional processing, detecting subtle links between physiological reactivity, social disposition, and negative affect. This suggests that pupil dynamics provide a privileged window into the earliest, automatic stages of social-emotional processing, where bodily responses precede conscious recognition.

In conclusion, our work supports the idea of a continuous interplay between early physiological reactions and higher-order social cognition [62], supporting the view that embodied autonomic signals scaffold emotion recognition and guide social behaviour.

## Methods

### Participants

We collected data from sixty-eight participants (50 females, mean age of 25.4 years old, sd = 5.5). To explore correlations with a moderate effect size, with α = 0.05 and power = 0.8, we calculated an a priori minimum sample size of 67 participants, based on calculations performed in G*Power [63]. It should be noted that a 2×2×3 repeated-measures analysis of variance (ANOVA) requires a minimum sample size of 18 participants to attain a moderate effect size with α = 0.05 and power = 0.95. Inclusion criteria were being between 18 and 45 years old, having normal vision or corrected by contact lenses, being fluent in Italian, and having normal stereoacuity. The latter was assessed using the Frisby stereo-test (Clement Clarke International Ltd, Essex, UK) [64]. All participants were able to detect the target on the thickest plate of the Frisby test from a distance of 60 cm, demonstrating a minimum stereoacuity of 150-sec arc. This level of stereo vision was deemed sufficient to ensure the dichoptic viewing required by the task. People with psychological or neurological diagnoses, or those with strabismus or amblyopia, were excluded from participation in the study. Fifteen participants had left-eye dominance, while 78% showed right-eye dominance. Participants were primarily recruited through announcements on major social media platforms and public advertisement boards in highly frequented spaces (a library, a university, etc.). A compensation of ten euros was given for 90 minutes of experimental participation. Some participants were recruited via the online university platform Sona Systems and were compensated with 90 course credits. Written consent was obtained from all participants prior to testing. Procedures were approved by the research ethics committee of the University of Trento. The study was performed in accordance with Helsinki’s declaration.

### Apparatus

The task was programmed and presented using E-prime 2.0 (Psychology Software Tools, Pittsburgh, PA, USA) on an Eizo FG2421 LCD Monitor with a resolution of 1920 x 1080 pixels and a refresh rate of 240 Hz. A stereoscopic mirror and a wood divider were placed in front of the screen to ensure a dichoptic view of the stimuli. A chin rest was placed under the stereoscopic mirror at a distance of 60 cm from the screen. Facial EMG activity and EDA were recorded from an MP160 Biopac modular system (Biopac Systems, Goleta, CA) through the software Acknowledge (Biopac Systems, Goleta, CA). Pupil diameter was measured using Tobii 2 eye-tracking glasses.

### Stimuli

Facial stimuli were selected from the FACES database of emotional expressions [65]. Fearful, happy, and neutral expressions were chosen from twelve different young individuals of matched gender (six females and six males). Images were edited using the GNU image manipulation program (GIMP). A grayscale version of the images was created, with the original background replaced by a uniform gray background (luminance = 25.14 cd/m^2^). A Konica Minolta LS-100 luminance photometer was used to measure luminance levels. The luminance of the stimuli was adjusted to correspond to a mean value of 21.18 cd/m^2^. A luminance-matched scrambled version of each image was created, while preserving the power spectrum and root mean square contrast [66]. The final set consisted of 72 images, divided into six conditions (3 emotions x 2 image types), with 12 images in each condition. An example of the stimuli is represented in Figure 7a. To control for potential effects of low-level visual features, we measured the spatial frequencies of the stimuli and compared the percentages of low-level frequencies across stimulus categories (results can be found in the Supplementary Materials). The faces and scrambled images were presented on the left side of the screen 500 ms after the beginning of the trial. Target contrast ramped up progressively from 0 to 100% over a period of one second to avoid abrupt presentation and remained constant at 100% until the end of the trial (see Figure 7b).

**Figure 7.**
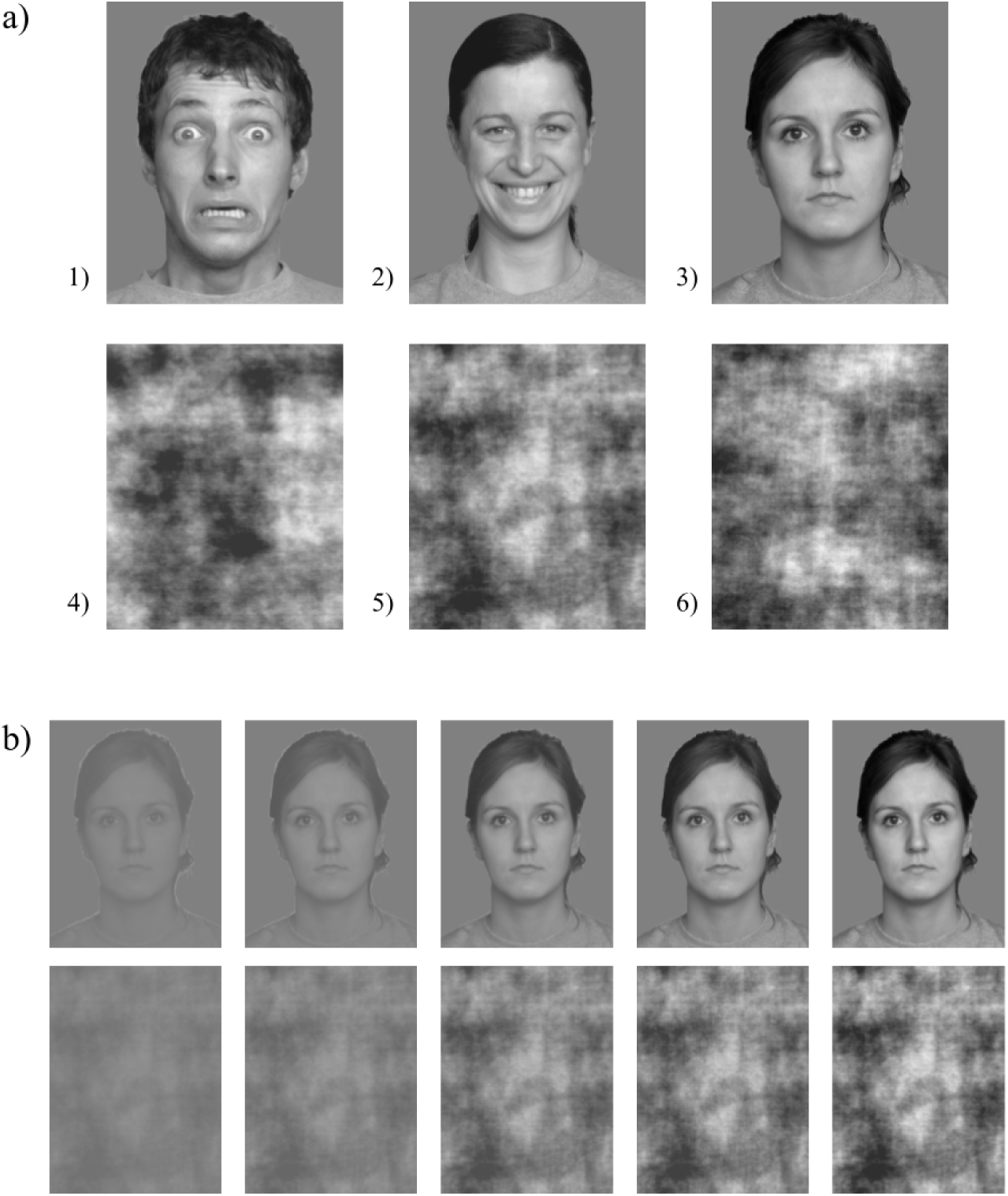
**a)** Examples of visual stimuli (selected from the FACES database of emotional expressions [65]) for each experimental condition: fearful (1), happy (2), neutral (3) faces in the top row, with the corresponding scrambled images in the bottom row (4-6). **b)** The ramp-up procedure. The figure illustrates the five different levels of contrast in which each stimulus was shown, ranging from 0 % up to 100 %, for faces (top) and scrambled images (bottom).

A flickering colored mask was displayed on the right side of the screen. The mask consisted of five distinct high-contrast Mondrian patterns, which cycled in sequential order at a refresh rate of 20 Hz (1 image every 50 ms) (e.g., [67]). A checkered frame (see Figure 8) composed of smaller grayscale squares surrounded both stimuli on either side of the divider to aid binocular fusion, while a small white fixation cross was placed in the middle of both stimuli to promote stable fixation. Both static and dynamic stimuli were preceded by a two-second gray fixation cross on a white background set to maximum luminance of 124,92 cd/m2.

**Figure 8.**
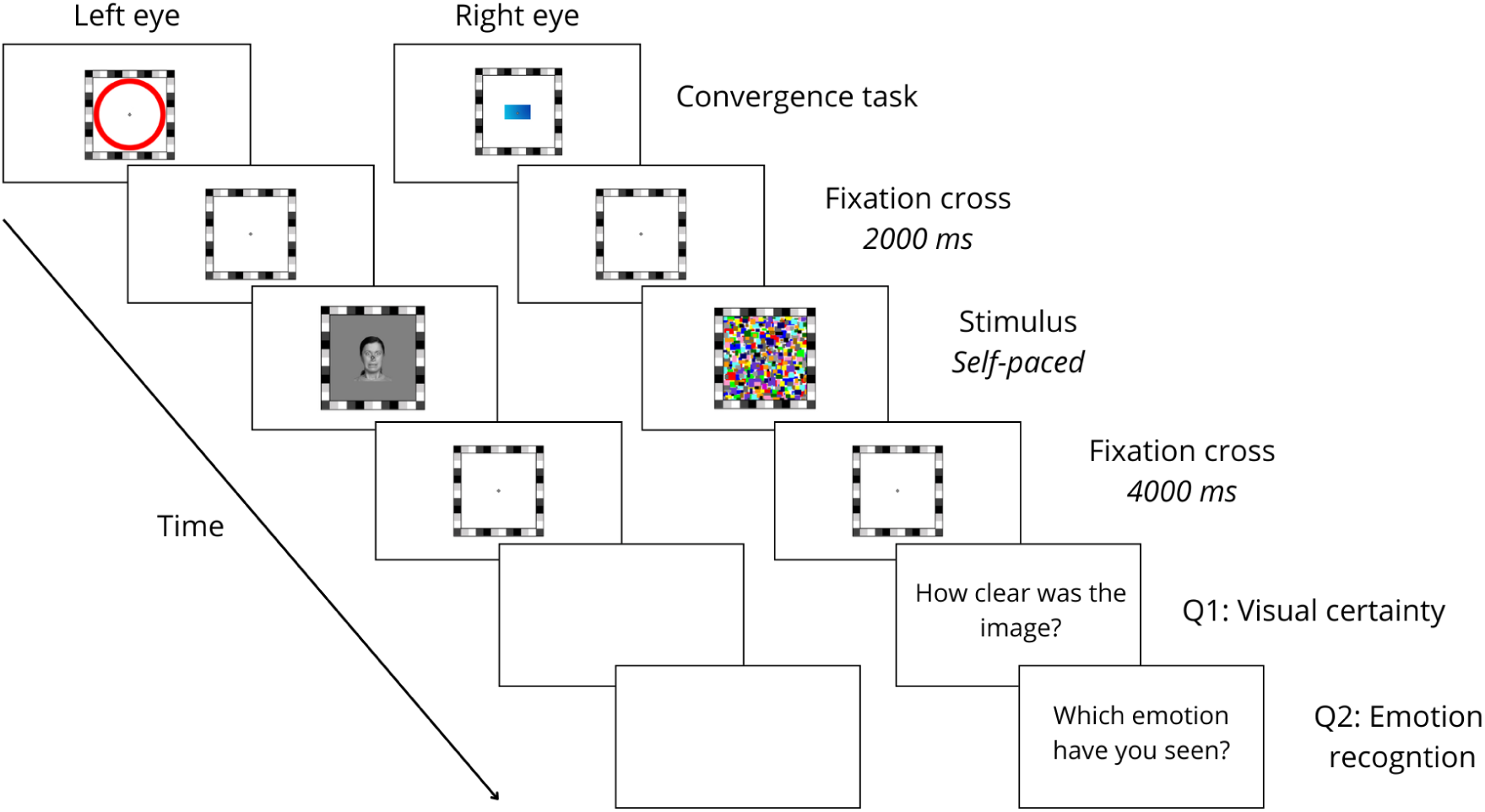
Trial sequence. Each stimulus could last up to a maximum of 10,500 ms. Trial duration was approximately 15 seconds on average.

### Procedure

After obtaining informed consent from each participant, we assessed their stereopsis using the plates of the Frisby stereopsis test [64] and annotated their ocular dominance. Participants were then prepared for the physiological data acquisition: electrodes for EDA and facial EMG recordings were placed and they were asked to wear Tobii 2 eye-tracking glasses (see the measurements section for more details). Physiological measurements were recorded throughout the entire experiment. Participants were seated with their chins fixed on a chin rest in a dimly lit room, and they were allowed to adjust the height of the chair for comfort. A brief demo of the experiment was conducted to ensure that participants understood the instructions and could correctly fuse the two images into one coherent percept. Participants were asked to press the spacebar as soon as they saw something different from the flickering mask. We emphasized the importance of pressing the button immediately, without taking time to further check the appearing image. All trials began with a convergence task to ensure that binocular fusion was achieved and maintained throughout the whole experiment. In this task, a red circle was presented to the left eye and a blue rectangle to the right eye. Participants were instructed to press the spacebar to commence the trial only after they perceived the rectangle in the middle of the circle as a single, coherent image. Then, a pre-stimulus fixation cross was presented for two seconds, followed by the simultaneous presentation of the static image to the left eye and the dynamic flickering mask to the right eye. These stimuli were presented until participants pressed the button to indicate the breakthrough of the target or after a maximum duration of 10,500 ms. A post-stimulus fixation cross was then presented for four seconds. At the end of the trial, participants were asked to respond to two forced-choice questions: one regarding visual certainty and the other one on emotion recognition. Emotion recognition was assessed by asking participants to choose among four different emotions, indicating which one they perceived if they saw a face, or stating that they did not see a face if none was perceived. Participants indicated their responses by showing the corresponding number with their fingers, minimizing unnecessary movements, and the researcher recorded the answer by pressing corresponding keys on the keyboard. Visual certainty was assessed using an adapted version of the perception awareness scale with a four-point Likert scale ranging from “I have seen nothing” to “The stimulus was completely clear” (PAS [21]). A schematic representation of the trial design is presented in Figure 8. The bCFS experiment lasted between 20 and 40 minutes, depending on the participant’s speed. Finally, participants were asked to complete the last set of questionnaires (see Questionnaires section). The total duration of the laboratory session was approximately 1 hour and 10 minutes.

### Questionnaires

To measure personality traits and emotional states, participants completed six questionnaires, which were divided into two sets. The first set of questionnaires included a brief demographic survey, the Autism Spectrum Quotient (AQ [68]), the short version of the Empathy Quotient (EQ [69]), and the Toronto Alexithymia Scale (TAS [70]).

In brief, the AQ is a self-report measure of autistic traits consisting of 50 items that can be grouped into five subscales: attention to detail, attention switching, imagination, communication, and social skills. Higher scores on the AQ correspond to higher levels of autistic traits. We adopted an Italian version, which was validated by Ruta and colleagues [71]. The EQ-40 is a self-report tool that assesses empathy in adults. It consists of 40 items divided into three subscales: cognitive empathy, emotional empathy, and social skills. It provides a total score that ranges between 0 and 80, with higher scores indicating higher levels of empathy. We administered the Italian version validated by Preti and colleagues [72]. The TAS-20 measures difficulty in identifying and describing emotions, which are both core characteristics of alexithymia. The 20 items can be categorized into three different components of alexithymia: difficulty identifying feelings, difficulty describing feelings, and externally oriented thinking. Higher scores indicate higher levels of alexithymic traits. The Italian version of the TAS-20 that we used was validated by Bressi and colleagues [73].

The second set of questionnaires included the Depression Anxiety Stress Scale (DASS-21 [74]), the Systemizing Quotient (SQ [75]), and the Schizotypal Personality Questionnaire - Brief (SPQ-BR [76]). The DASS-21 consists of 21 questions measuring the levels of Depression, Anxiety, and Stress experienced in the last week. We administered the Italian version of the DASS validated by Bottesi and colleagues [77]. The SQ is used to assess systemizing cognitive style, which is the tendency to classify and categorize objects and information. This 75-item self-report questionnaire is widely used to complement other measures of autistic traits, as autistic individuals tend to score high on this cognitive dimension. We used the Italian version of the SQ validated by Ruta et al. (2003). Finally, the SPQ-BR (32 items) assesses schizotypy levels. The SPQ-BR includes seven different subscales: suspiciousness, magical thinking, unusual perceptions, social anxiety, constricted affect, eccentric behaviour, and odd speech. Higher scores in the global or subordinate scales correspond to higher levels of schizotypal traits. The Italian version has been validated by Fossati and colleagues [78].

All questionnaires were administered through the online platform Qualtrics. The first set of questionnaires was sent to participants before the lab experiment, to be completed at home, while the second set was filled out in the lab after the bCFS experiment.

### Measurements

#### Electrodermal activity

Before applying the electrodes to the middle and index fingers of the left hand, we asked participants to clean the area with a disinfectant wipe and then dry it carefully. Two dry, reusable Silver-Silver Chloride (Ag-AgCl) finger electrodes were filled with isotonic gel and strapped to the distal phalanges of the left middle and index fingers. The signal was recorded using the EDA 100C BIOPAC Systems module (2000 Hz sampling rate, Gain: 5 μV, 10 Hz low-pass filter), and triggers were sent from the presentation software via a parallel port. The signal was pre-processed offline using Matlab’s Ledalab toolbox [79]. The signal was down-sampled to 10Hz, and a Butterworth low-pass filter was applied. It was then decomposed into tonic and phasic components by means of continuous decomposition analysis, and finally, the mean values for the window of interest (1 to 3 seconds after the event) were extracted. Two types of events were considered for different analyses: the stimulus onset and the participant’s response.

Six participants were excluded from the analysis due to corrupted data. Eleven additional participants were excluded because they were considered as outliers, with values exceeding 2.5 SD of the mean, resulting in a final sample of 51 valid participants for the analysis.

#### Facial EMG

To improve adherence of the electrodes and remove traces of dirt or dead skin, we used Nuprep cream to scrub the left cheek (i.e., area of the zygomaticus major) and the left eyebrow area (i.e., corrugator supercilii), where electrodes were then placed according to the guidelines by Fridlund and Cacioppo [80]. Two adjacent electrodes were placed on each muscle of interest, and the ground signal was obtained from the EDA electrodes, following the guidelines provided by the Biopac producers. These muscles were selected based on extensive literature showing greater activation of the zygomaticus in response to smiling faces and greater activation of the corrugator in response to frowning (angry or fearful) faces. The muscular activity was amplified through a Biopac MP160 (Biopac Systems, Goleta, CA) and recorded with the associated AcqKnowledge software. The signal was online-filtered with a 500 Hz low-pass filter and a 10 Hz high-pass filter. The sampling rate was set to 2000 Hz, and a notch filter at 50 Hz was applied to reduce line noise. EMG offline pre-processing started with a bandpass filter of 30-300 Hz. The filtered signal was then rectified, integrated, and cut into epochs. We created two different time windows, one locked to the stimulus onset (from 0 to 5s) and the other one locked to the response time (from −1s to 5s). For each time window, data were normalized by baseline division. Specifically, the mean EMG value from the 500 ms preceding stimulus onset was used as the baseline for stimulus-locked analysis, while the mean EMG value from a 500 ms interval occurring 3 seconds after the response (during the post-stimulus fixation cross) was used for response-locked analysis. Dimberg and Thunberg (1998) [30] showed that muscle reactions to facial expressions occur within 500 ms after the stimulus onset. Following this finding, mean amplitudes of the first 500 ms bin were extracted and averaged across conditions for statistical analysis.

Seven participants were excluded from the analyses due to corrupted data. Six additional participants were excluded because they were identified as outliers (values exceeding 2.5 SD of the mean), resulting in a final sample of 55 valid participants for the analysis.

#### Pupillometry

Pupil diameter was measured using the Tobii Pro Glasses 2 with a sampling rate of 100 Hz. Participants’ gaze was calibrated using a bullseye card, which they were asked to hold at arm’s length. Pupil data was pre-processed and analyzed using the CHAP software [28].

Eye blinks were detected using the algorithm developed by Hershman and colleagues [81], and the missing data points were replaced using a linear interpolation method [82]. Following recommended practices in pupillometry preprocessing (e.g., *CHAP* pipeline and field-standard thresholds [28,83]), sample-level outliers exceeding ± 2.5 z-scores were removed. Trials with more than 25% missing or invalid data were excluded from analysis (consistent with thresholds tested by Burg and colleagues [84]). Participants retained only if at least 80% of trials per condition (minimum of 10 trials) remained valid, ensuring sufficient within-subject reliability. Finally, we created two sets of time-course data points by aligning the onset of the time window either to the stimulus onset or to the response time (i.e., the moment when the participant noticed something breaking through the suppression). In both cases, the data were normalized to their baseline, defined as the average pupil size 500 ms prior to the time window of interest.

Fifteen participants were excluded from the analysis due to corrupted data, and three additional participants were excluded because they did not reach the threshold of valid trials, resulting in a final sample of 50 valid participants for the pupil analysis.

### Data analysis

A two-way repeated measures ANOVA was carried out on mean RTs and accuracy percentage in emotion recognition with Image Type (intact vs. scrambled) and Emotion (fearful vs. happy vs. neutral) as the main factors. For the analysis of the EMG and EDA data, a third factor assessing the effect of Awareness was added to the ANOVA, resulting in a three-way repeated-measures ANOVA testing Image Type, Emotion, and Awareness (pre vs. post) as main factors. Any significant interactions were followed up with post-hoc comparisons, using Bonferroni correction. To analyze any differences in pupil dilation across conditions, we followed Hershman and colleagues [28] approach, which consisted of running a series of Bayesian paired-samples t-tests between conditions over the averaged time course of the two epochs that corresponded to the pre-(stimulus-locked) and post-(response-locked) awareness phases. We first compared the mean dilation for faces versus the mean dilation for scrambled images, and then separately tested the three emotions of the intact faces and their scrambled versions.

We further controlled for potential influences of perceptual awareness by correlating the PAS scores with the behavioural (RTs) and physiological measures. More specifically, we adopted the Spearman’s rank correlation coefficient, given the non-normal distribution of the PAS scores. No significant correlations were found (all ps > .05); therefore, we excluded any potential effects of varying degrees of perceptual awareness in the processing of the stimuli.

To examine the influence of interindividual differences on behavioural and physiological responses, one-tailed bivariate correlations were conducted between questionnaire scores and arousal indices, in line with our a priori directional hypotheses. Spearman’s rank correlation coefficients (*rs*) were used for non-normally distributed data, whereas Pearson’s product–moment correlation coefficients (*r*) were employed otherwise.

To reduce the number of comparisons, we calculated a single value of social stimuli sensitivity for all indices by subtracting the mean values of the scrambled images from the mean values of the intact faces. In doing so, we obtained two values for each physiological index, one before and one after visual awareness. For this analysis, the mean pupil dilation was calculated over a 500 ms time window starting 1,500 ms after stimulus onset or after the response. The significant correlations were further explored using multiple linear regression (forward selection) to identify which of the different questionnaires’ subscales best predicted the physiological response. All the analyses were performed in SPSS (version 25, IBM Corporation, Armonk, NY).

## Acknowledgments

We wish to thank V. Dapor for constructing the apparatus, P. Leoni for providing technical support, A. Chiavassa, F. Cestaro, A. Zuriati, G. Marcolin, and A. Tomaselli for helping with data collection.

## Data and code availability

Data for the reported experiment is currently available via the Open Science Framework (OSF): https://osf.io/zeujg/?view_only=cdc8bd7e958849b7bfcd3f629dbe31df. The Matlab codes used in the analysis of this study are available on request from the author C.D.

## Authors contribution

C.D.: conceptualization, data curation, formal analysis, investigation, methodology, writing - original draft; R.H: formal analysis, writing - review and editing; D.R: formal analysis, writing - review and editing; F.M.: conceptualization, data curation, funding acquisition, methodology, supervision, writing - original draft; I.S.: conceptualization, data curation, funding acquisition, methodology, supervision, writing - original draft. All authors gave final approval for publication and agreed to be held accountable for the work performed therein.

