## Supplementary Materials for "Pre-conscious reactions to faces as the biological roots of emotion perception"

### Appendix A. Spatial frequencies analysis

To rule out any potential effects of low-level visual features, we performed a Spatial Frequency analysis using the Natural Image Statistical Toolbox [1]. Specifically, we employed the toolbox script to compute the spectral energy distribution across our image set. While maintaining the default cumulative energy level at 80%, we adjusted the cycle-per-image threshold to 3 cycles/image to focus on low spatial frequencies. We then calculated the proportion of spectral energy falling below this threshold. We focused specifically on low spatial frequencies, as the subcortical visual pathway primarily processes these coarse frequencies [2]. Repeated-measures ANOVAs were conducted on the low spatial frequencies percentages with Image Type (Intact vs. scrambled) and Emotion (Fearful vs. Happy vs. Neutral) as main factors.

The repeated-measures ANOVA revealed a significant main effect of Image Type ($F_{1,11}$ = 5.35, p = .041, ηp2 = .327), with intact images having smaller percentages of low frequencies than scrambles. The main factor Emotion ($F_{2,22}$ = 2.33, p = .121, ηp2 = .17) and the interaction between Image Type and Emotion ($F_{2,22}$ = 1.25, p = .305, ηp2 = .102) did not reach significance (Figure A).

It is highly unlikely that the slightly higher percentages of low spatial frequencies in the scrambled images, compared to the intact faces, accounted for the observed effects on behavioural performance. If this difference had been influential, we would have expected shorter reaction times (RTs) for the scrambled images; however, our findings revealed the opposite pattern. Overall, the analysis of the low spatial frequencies indicates that differences in RTs were not driven by low-level image features.

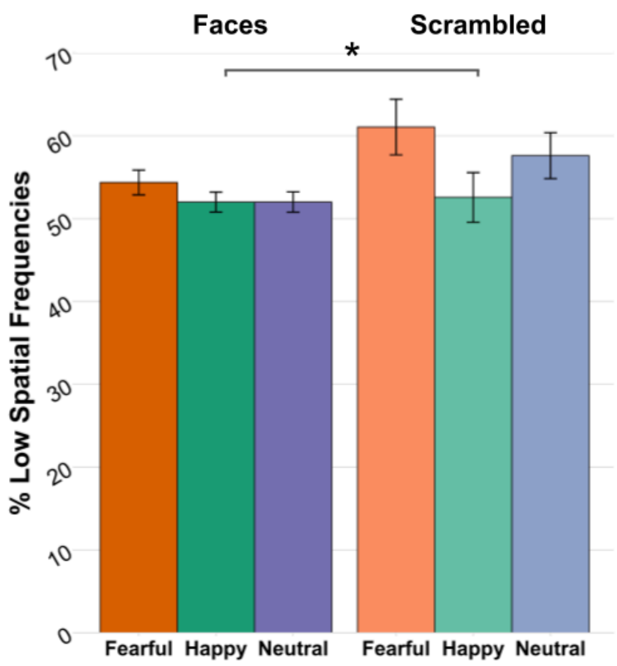


***Figure A.*** *Mean percentages of low spatial frequencies for all the conditions. Error bars correspond to standard error of the mean. Asterisks denote significant differences at p < .05 (*).*

### Appendix B. Perceptual awareness

To assess whether perceptual awareness influenced physiological responses, we examined participants' ratings on the Perceptual Awareness Scale (PAS) [1]. Due to the uneven distribution of PAS scores within and across conditions, other types of trial-level analysis were not feasible. Instead, we averaged PAS scores for each condition and correlated them with the corresponding values of each physiological index. None of the correlations reached significance (*r*s ranged from −.25 to .25, all *ps* > .05, one-tailed), indicating that pre-conscious physiological responses were not modulated by participants' subjective awareness.

### Appendix C. Correlations analysis

***Correlations between personality traits and physiological responses after visual awareness***

Correlations between personality traits, emotional states, and physiological indices were also computed for the conscious phase. As it turned out, DASS correlated positively with the activation of both the zygomaticus ($rs(53) = .259$, $p_{uncorr} = .03$) and the corrugator ($rs(54) = .362$, $p_{uncorr} < .01$) muscles. A multiple regression was conducted to examine whether DASS total, DASS Depression, and DASS Anxiety scores predicted EMG responses to facial stimuli. The overall model was significant, $F(3, 51) = 3.57$, $p = .020$, and accounted for approximately 17.3% of the variance in EMG responses, $R^2 = .17$, adjusted $R^2 = .13$. Among the predictors, only DASS Anxiety was a significant predictor of EMG responses, $\beta = .64$, $t = 2.43$, $p = .019$. Neither DASS Depression ($\beta = .07$, $p = .76$) nor DASS total score ($\beta = –.33$, $p = .35$) were significant. DASS Stress was excluded from the model due to multicollinearity (tolerance = .000). Given the positive trend observed in both muscles, we interpret this result not as a mimicry effect, but rather as a general muscular tension expressed in the face. This suggests that higher levels of anxiety are associated with a “nervous” facial reaction to faces, consistent with previous findings in the literature [1–3].

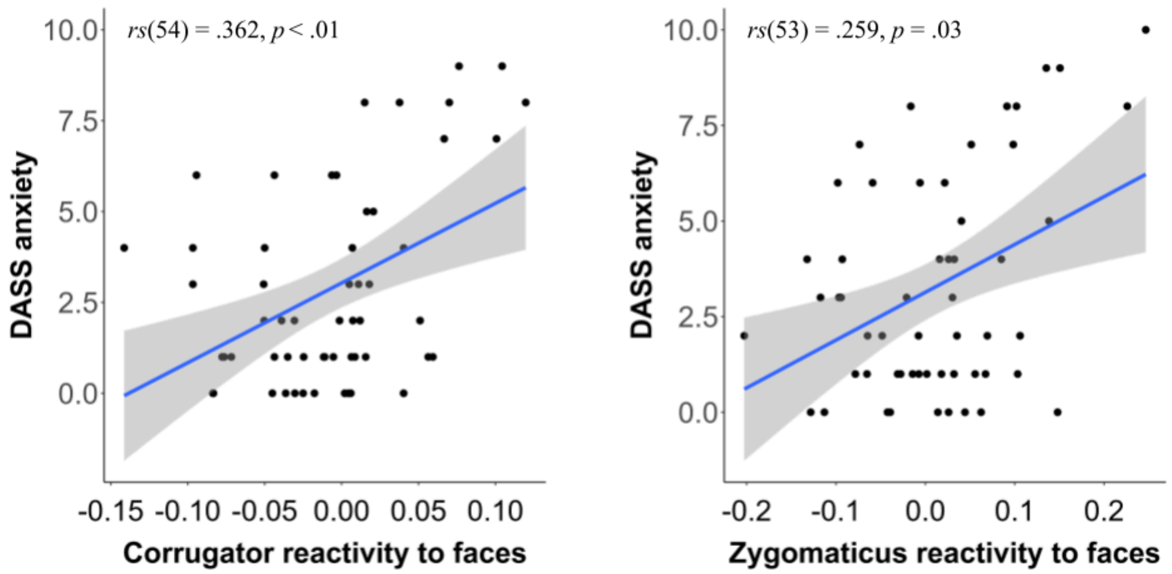


***Figure C.*** *Correlations between conscious EMG activation and DASS anxiety scores. Positive values on the x axis indicate greater activity toward faces compared to scrambled images.*

### Appendix D.

### Experiment 2. Detection task.

To rule out any potential effects due to task demands, we carried out a control experiment, where the emotion recognition task was replaced by a simple detection task. It is possible that the emotion recognition task used in the main experiment might have slowed down participants' RTs. In particular, participants may have retarded their button press to ensure accurate recognition of the emotion expressed by the face. In this control experiment, we expected to observe faster RTs; nonetheless, we still anticipated a privileged access to consciousness for social stimuli.

#### Methods

***Participants***

Nineteen participants (11 females, mean age of 25.5y, sd = 5.9) took part in Experiment 2. The inclusion and exclusion criteria were identical to those described in the main experiment. Participants were mainly recruited through announcements on the major social media platforms and public advertisement boards in highly frequented spaces (a library, a university, ecc). A compensation of ten euros was given for 90 minutes of experimental participation. Some participants were recruited via the online university platform Sona, and were compensated with 90 course credits. Written consent was obtained prior to testing and all procedures were approved by the Research Ethics Committee of the University of Trento. The study was performed in accordance with the Declaration of Helsinki.

**Stimuli and procedure**

The same stimuli and procedure described for experiment 1 were used. The only modification was the removal of the emotion recognition question. After stimulus detection, participants only responded to the visual certainty scale. We focused on the behavioural data, analysing only the breakthrough times. For the statistical analysis, we followed the same approach used in the main study and performed a 2x3 repeated-measures ANOVA on RTs with Image Type (intact vs. scrambled) and Emotion (fearful vs. happy vs. neutral) as main factors.

### Results experiment 2

#### *Behavioural results*

Similarly to experiment 1, the mean RT for faces (mean = 1914, SD = 856) was significantly faster than the mean RT for the scrambles (mean = 2993, SD = 1869) (i.e., main effect of Image Type: $F_{1,18} = 16.902$, $p < .01$, $\eta p^2 = .484$). The analysis also revealed a significant interaction between ImageType and Emotion ($F_{2,36} = 3.5$, $p = .04$, $\eta p^2 = .163$). Post-hoc comparisons only revealed a trend showing that neutral expressions broke through suppression faster than happy expressions ($p = .055$). No significant effect was found for Emotion ($F_{2,36} = .22$, $p = .80$).

Overall, the data suggest that modifying the experimental task altered the pattern of RTs. Specifically, reaction times were generally shorter in this detection task compared to Experiment 1, likely due to reduced processing demands; in Experiment 1, participants were required to identify emotional expressions and provide a response, which may have prolonged RTs. Unlike in Experiment 1, happy expressions were no longer detected the fastest; instead, neutral expressions showed the shortest RTs.

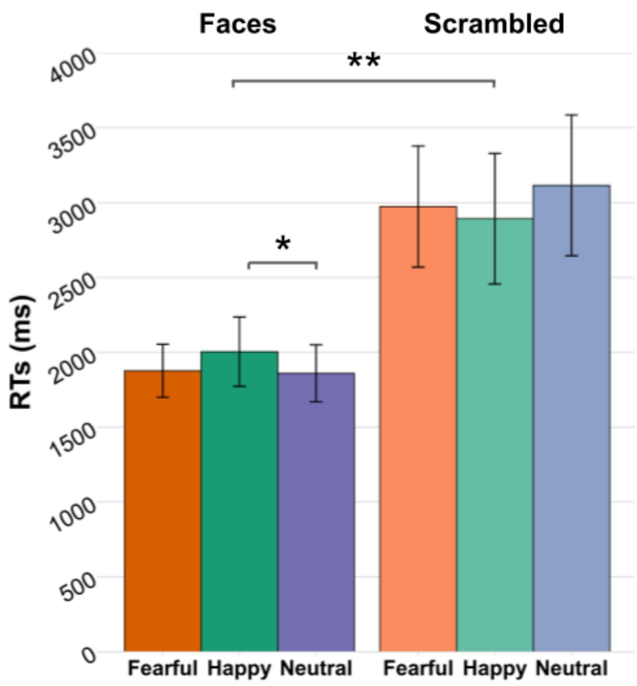


***Figure D.*** *Mean breakthrough times for all the conditions. Error bars correspond to standard error of the mean. Asterisks denote significant differences at p < .05 (*) and p < .01 (**).*
